# Reverse-Complement Equivariant Networks for DNA Sequences

**DOI:** 10.1101/2021.06.03.446953

**Authors:** Vincent Mallet, Jean-Philippe Vert

## Abstract

As DNA sequencing technologies keep improving in scale and cost, there is a growing need to develop machine learning models to analyze DNA sequences, e.g., to decipher regulatory signals from DNA fragments bound by a particular protein of interest. As a double helix made of two complementary strands, a DNA fragment can be sequenced as two equivalent, so-called *Reverse Complement* (RC) sequences of nucleotides. To take into account this inherent symmetry of the data in machine learning models can facilitate learning. In this sense, several authors have recently proposed particular RC-equivariant convolutional neural networks (CNNs). However, it remains unknown whether other RC-equivariant architectures exist, which could potentially increase the set of basic models adapted to DNA sequences for practitioners. Here, we close this gap by characterizing the set of all linear RC-equivariant layers, and show in particular that new architectures exist beyond the ones already explored. We further discuss RC-equivariant pointwise nonlinearities adapted to different architectures, as well as RC-equivariant embeddings of *k*-mers as an alternative to one-hot encoding of nucleotides. We show experimentally that the new architectures can outperform existing ones.

## 1 Introduction

Incorporating prior knowledge about the structure of data in the architecture of neural networks is a promising approach to design expressive models with good generalization properties. In particular, exploiting natural symmetries in the data can lead to models with fewer parameters to estimate than agnostic approaches. This is especially beneficial when the amount of available data is limited. A famous example of such an architecture is the convolutional neural network (CNN) for 1D sequences or 2D images, which is well adapted to problems which are invariant to translations in the data, while exploiting multiscale and local information in the signals. Motivated by the success of CNNs, there has been a fast-growing body of research in recent years to build the theoretical underpinnings and design architectures and efficient algorithms to systematically exploit symmetries and structures in the data [3].

A central idea that has emerged is to formalize the symmetries in data by a particular *group action* (e.g., the group of translations or rotations on images), and to create multilayer neural networks which, by design, “behave well” under the action of the group. This is captured formally by the concept of *equivariance*, which states that each equivariant layer should be designed to be subject to the group action (e.g., we should be able to “translate” or “rotate” the signal in each layer), and that when an input data is transformed by a particular group element, then its representation in an equivariant layer should also be transformed according to the same group element. While it is easy to see that convolutional layers in CNNs are equivariant to translations, Cohen and Welling [7] formalized the concept of group equivariance CNN (G-CNN) for more general groups and showed in particular how to design convolutional layers equivariant not only to translations but also to reflections and to a discrete set of rotations. Following this seminal work, the theoretical foundations of group equivariant neural networks were then expanded, going beyond regular representations [9], for more groups [2, 18, 37, 40], in less regular spaces [8, 10] or with more general results on their generality and universality [11, 13, 14, 22]. The main applications were developed with the groups of rotations in 2D and 3D, mostly to computer vision problems, but also in biology with histopathology [17, 23], medicine [41] and quantum chemistry [33].

In this paper, we explore and study the potential benefits of equivariant architectures for an important class of data, namely deoxyribonucleic acid (DNA) sequences. DNA is the major form of genetic material in most organisms, from bacteria to mammals, which encodes in particular all proteins that a cell can produce and which is transmitted from generation to generation. The study of DNA in humans and various organisms has led to tremendous progress in biology and medicine since the 1970s, when the first DNA sequencing technologies were invented, and the collapsing cost of sequencing in the last twenty years has accelerated the production of DNA sequences: there are for example about 2.8 billion sequences for a total length of ~ 10^13^ nucleotides publicly available at the European Nucleotide Archive (ENA^1^). Unsurprisingly in such a data-rich field, machine learning-based approaches are increasingly used to analyze DNA sequences, e.g., in metagenomics to automatically predict the species present in an environment from randomly sequenced DNA fragments [26, 28, 36, 38] and to detect the presence of viral DNA in human samples [36], in functional genomics to predict the presence of protein binding sites or other regulatory elements in DNA sequences of interest [16, 24, 30, 35, 43, 44], to predict epigenetic modifications [25], or to predict the effect of variations in the DNA sequence on a phenotype of interest [1, 46].

Due to the sequential nature of DNA and the translation-equivariant nature of the questions addressed, many of these works are based on 1D CNN architectures, although recently transformer-based language models have also shown promising results on various tasks [6, 20, 42]. However, besides translation, DNA has an additional fundamental symmetry that has been largely ignored so far: the so-called reverse complement (RC) symmetry, due to the fact that DNA is made of two strands oriented in opposite direction and encoding complementary nucleotides. In other words, a given DNA segment can be sequenced as two RC DNA sequences, depending on which strand is sequenced; any predictive model for, e.g., DNA sequence classification should therefore be RC-invariant, which calls for RC-equivariant architectures. While strategies based on data augmentation and prediction averaging has been commonly used to handle the need for RC invariance [1, 32], one translation- and RC-equivariant CNN architecture has been proposed and led to promising results [4, 29, 34]. However, it remains unclear whether that architecture is the only one that can encode translation- and RC-equivariance, or if alternative models exist to complement the toolbox of users wishing to develop deep learning models for DNA sequences.

Using the general theory of equivariant representations, in particular steerable CNNs [9], we answer that question by characterizing the set of all linear translation- and RC-equivariant layers. We show in particular that new architectures exist beyond the ones already explored by [4, 29, 34], which in the language of equivariant CNNs only make use of the regular representation [7] while more general representations lead to different layers. We further discuss RC-equivariant pointwise nonlinearities adapted to different representations, as well as RC-equivariant embeddings of *k*-mers as an alternative to one-hot encoding of nucleotides. We test the new architecture on several protein binding prediction problems, and show experimentally that the new models can outperform existing ones, confirming the potential benefit of exploring the full set of RC-equivariant layers when manipulating DNA sequences with deep neural networks.

## 2 Methods

### 2.1 Group action of translation and reverse complementarity on DNA sequence

DNA is a long polymer made of two intertwined strands, forming the well-known double-helical structure. Each strand is a non-symmetric polymer that can be described as an oriented chain of four possible monomers called nucleotides and denoted respectively {`A, C, G, T`}. The two strands are oriented in opposite directions, and their nucleotides face each other to form hydrogen bonds. They interact at each position in a deterministic way because only two nucleotides pairings can happen: (`A,T`) and (`G,C`). Thus, given a nucleotide sequence on one strand, we can deduce the so-called RC sequence of its corresponding strand by complementing each nucleotide and reversing the order (**Figure 1)**. When a double-stranded DNA fragment is sequenced, the two strands are first separated and, typically, only one of them is randomly selected and is decrypted by the machine. This implies that any given DNA fragment can be equivalently described by two RC sequences of nucleotides. Moreover, several genomic learning tasks amount to a sequence annotation that does not depend on the strand. For example, a protein can bind a double-stranded DNA fragment, and both strands of the bound part can get sequenced. This motivates the search for equivariance to this RC-action for the prediction functions. Moreover, the sequencing often results in long sequences where the relevant parts of the sequence do not correlate with their position. The task of prediction over genomic sequences is thus largely translation equivariant, which explains why the community settled on the use of CNNs to train and predict on arbitrary length segments.

**Figure 1:**
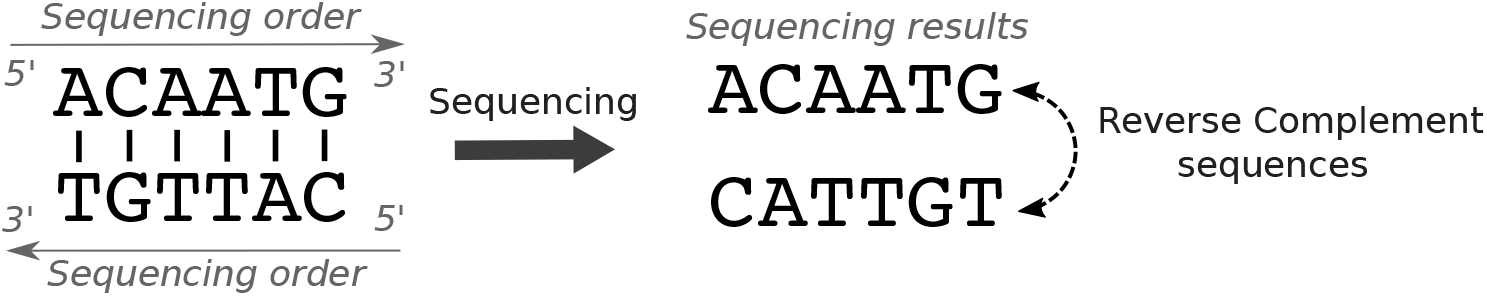
Illustration of the reverse-complement symmetry. Both DNA strands get sequenced in opposite directions resulting in redundant information.

To formalize mathematically the translation and RC operations on DNA sequences, we first encode the raw genetic sequence as a signal function in 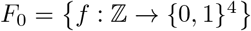, as the one hot encoding of the nucleotide content for each integer position. Because of the finite length of this polymer, we assume that beyond a compact support this function takes a constant value of zero. The group 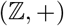 of translations acts naturally on this encoding by *T_u_*(*f*)(*x*) = *f*(*x − u*), for a translation 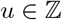, and the RC operations amounts to the following : *RC*(*f*)(*x*) = *σ*(−1)[*f*(−*x*)], where *σ*(−1) is the 4 × 4 permutation matrix that exchanges complementary bases `A/T` and `C/G` (while we denote by *σ*(1) the 4 × 4 identity matrix). We notice that *RC* is a linear operation on *F*_0_ that satisfies *RC*^2^ = *I*, and thus that the RC operation is a group representation on *F*_0_ for the group 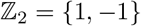 endowed with multiplication.

To jointly consider translations and RC actions, we naturally consider the semi-direct product group 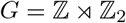. Elements *g* ∈ *G* can be written as *g* = *ts* with 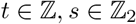 and the group *G* acts on *F*_0_ by the action *π*_0_ defined by:

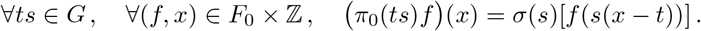

In other words, *π*_0_ is the representation of *G* on *F*_0_ induced by the representation *σ* of RC on 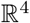 [9].

### 2.2 Features spaces of equivariant layers

Let us now describe the structure of intermediate layers of a neural network equivariant to translations and RC. Following the theory of steerable CNNs [9], we consider successive representations of the input DNA sequence in the following way:

#### Definition 1.

*Given ρ a representation of* 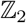 *on* 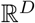 *for some* 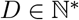, *a ρ-feature space is the set of signals* 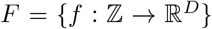 *endowed with the G group action π, known as the representation induced by ρ*:

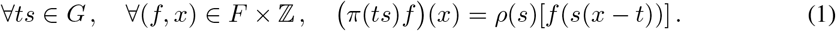

With this definition, we see in particular that the one-hot encoding input layer maps the input DNA sequence to a *σ*-feature space, and that the dimension (i.e., number of channels in the language of deep learning) and group action of *ρ*-feature space are fully characterized by the representation *ρ*. Interestingly, the theory of linear group representations allows us to characterize more precisely *all* such representations:

#### Theorem 1.

*For any representation ρ of* 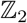 *on* 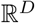, *there exist* 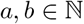 *such that a* + *b* = *D and an invertible matrix* 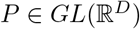 *such that*

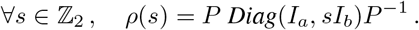

In other words, combining Definition 1 and Theorem 1, we see that any *ρ*-feature space that we will use to build translation- and RC-equivariant layers is fully characterized by a triplet (*P, a, b*), which we call its *type*, and which characterizes both its dimension *D* = *a* + *b* and the action of the group *G* by (1). By slight abuse of language, we also refer to (*P, a, b*) as the type of *ρ*.

Theorem 1 is a standard result of group theory, which explicits the decomposition of any representation *ρ* in terms of so-called irreducible representation, or *irreps*. In the case of 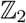, there are exactly two irreps which act on 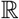, namely, *ρ*_1_(*s*) = 1 and *ρ*_−1_(*s*) = *s*. If *ρ* has type (*P, a, b*), then it means that it can be decomposed as *a* times *ρ*_1_(*s*) and *b* times *ρ*_−1_(*s*). In the particular case where *P* is the identity matrix, i.e., when we consider a type (*I, a, b*), then *ρ*(*s*) is a diagonal matrix for any 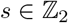, and each channel of *F* is acted upon by a single irrep. In that case, we will call the channels of type “1” (resp. “-1”) if they are acted upon by *ρ*_1_(resp. *ρ*_−1_), and we will say that *F* is an “irrep feature space”.

Now, let us introduce another special case. Since 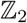 is finite of cardinality 2, let us consider the *regular representation ρ_reg_* of 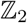 on 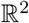 defined by:

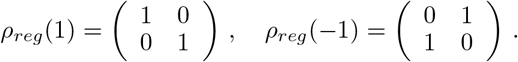

One can easily check that *ρ_reg_* is of type (*P_reg_*, 1, 1), where 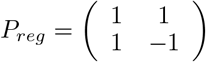. It corresponds to a *ρ*-feature space with two channels, where the RC operations flips the two channels (and of course the sequence coordinates).

Let us now consider feature spaces of interest. In the input layer, nucleotides are one-hot encoded in a certain order, let us say (`A, T, G, C`). As stated above, this input space is acted upon by *σ*, a 2–cycle that swaps bases `A/T` and `C/G`. We see that we can rewrite 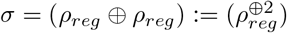, where ⊕ is the bloc-diagonal operation. Because *ρ_reg_* is of type (*P_reg_*, 1, 1), we can diagonalize *σ* with 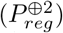 and the diagonal would be alternated +1 and −1 values. Thus, there exists a permutation Π such that *σ* is of type (*P*, 2, 2), with 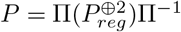. These concepts are illustrated in Supplementary Section A.1.

Interestingly, all RC-equivariant layers proposed so far in [4, 29, 34] follow a similar pattern: the channels go by pair, and the RC action amounts to flipping the channel values within a pair and reversing the sequence coordinates. In our formalism, this corresponds to channels of type (*P, a, a*), where 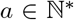 is the number of pairs of channels, and where up to a permutation of channels the matrix *P* satisfies 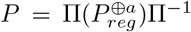. Following [34], we will refer to these layers as *Reverse Complement Parameter Sharing* (RCPS) layers below.

This highlights the fact that translation- and RC-equivariant layers explored so far are equivariant according to Definition 1, but that there exists potentially many other equivariant layers, obtained in particular by allowing *ρ*-feature spaces of types (*P, a, b*) where *a* ≠ *b*, on the one hand, and where *P* is not a direct sum of *P_reg_*, on the other hand. We investigate such variants below.

### 2.3 Equivariant linear layers

While Definition 1 characterizes *ρ*-feature space in terms of structure and group action, an equivariant multilayer neural network is built by stacking *ρ*-feature spaces on top of each other and connecting them with equivariant layers. Cohen et al. [11, Theorem 2] gives us a general result about such equivariant mappings. Here, we apply this result to our specific data and group, and characterize the class of equivariant linear layers, i.e., the linear functions *ϕ* : *F_n_* → *F*_*n*+1_ that satisfy *π*_*n*+1_*ϕ* = *ϕπ_n_*, where *π_n_* and *π*_*n*+1_ are respectively the group action on *F_n_* and *F*_*n*+1_.

#### Theorem 2.

*Given two representations ρ_n_ and ρ*_*n*+1_ *of* 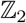, *of respective types* (*P_n_*, *a_n_, b_n_*) *and* (*P*_*n*+1_, *a*_*n*+1_, *b*_*n*+1_) *with a_n_* + *b_n_* = *D_n_ and a*_*n*+1_ + *b*_*n*+1_ = *D*_*n*+1_, *and respective ρ_n^-^_ and ρ*_*n*+1^-^_ *feature spaces F_n_ and F*_*n*+1_, *a linear map ϕ* : *F_n_* → *F*_*n*+1_ *is equivariant if and only if it can be written as a convolution*:

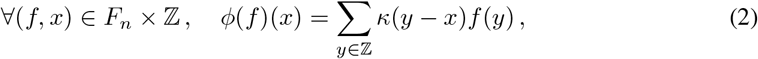

*where the kernel* 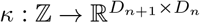 *satisfies*:

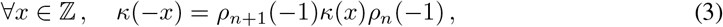

*or equivalently*:

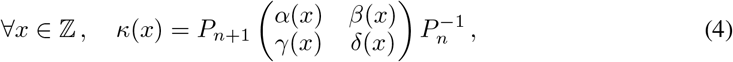

*where* 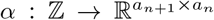 *and* 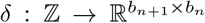 *are even, while* 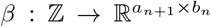 *and* 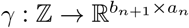 *are odd functions*.

As stated in Cohen et al. [11], “Convolution is all you need” to define linear layers which are equivariant to our group. In addition, Theorem 2 characterizes all the convolution kernels that ensure equivariance through the two equivalent constraints (3) and (4).

To illustrate this result, let us consider two RCPS feature spaces *F_n_* and *F*_*n*+1_ of respective types 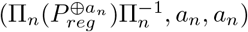 and 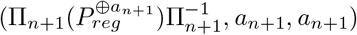. Then, the channels in *F_n_* and *F*_*n*+1_ go by pair, and if we consider a slice 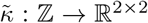 of the convolution kernel *κ* describing how a pair of channels in *F_n_* maps to a pair of channels in *F*_*n*+1_, (3) gives the constraint:

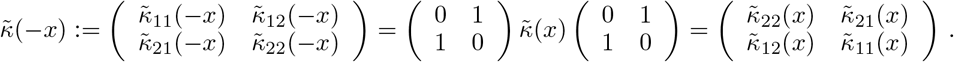

We recover exactly the constraints of the RCPS filters first proposed by [34], proving as a consequence of Theorem 2 that RCPS convolution filters describe exactly *all* equivariant linear mappings between RCPS feature spaces.

Moreover, if we now consider any two feature spaces *F_n_* and *F*_*n*+1_ of respective types (*P_n_, a_n_, b_n_*) and (*P*_*n*+1_, *a*_*n*+1_, *b*_*n*+1_), then Equation (4) tells us that up to multiplications by matrices *P*_*n*+1_ and 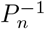, the kernel is expressed in terms of even and odd functions, which can be trivially implemented with parameter sharing. For example, to represent the even function *α*, one just need to parameterize the values of *α*(*x*) for *x* ≥ 0, and complete the negative values by parameter sharing *α*(−*x*) = *α*(*x*). Hence, the parameter sharing idea used in RCPS [34] extends to any equivariant linear map.

Instead of using (4) to parameterize equivariant convolution kernels, one may also directly write the constraints (3) for specific representations, and potentially save the need of multiplication by *P*_*n*+1_ and 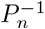 in (4). This is for example the case in RCPS layers [34], and more generally for channels acted upon by the regular representation; for the sake of completeness, we derive in Appendix A.4 the constraints to go from and to the regular representation or the irreps, and use them in our implementation.

### 2.4 Equivariant nonlinear layers

Besides equivariant linear layers, a crucial component needed for multilayer neural networks is the possibility to have equivariant nonlinear layers, such as nonlinear pointwise activation functions or batch normalization [19]. In this section, we discuss particular nonlinearities that are adapted to various equivariant layers.

#### Pointwise activations

Let us begin with pointwise transformations, that encompass most activation functions used in deep learning. Pointwise transformations are formally defined as follows: given a function 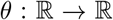 and a vector space 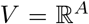 for some index set *A*, the pointwise extension of *θ* to *V* is the mapping 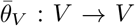 defined by 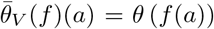, for any (*f, a*) ∈ *V* × *A*. For a *D*-dimensional representation *ρ* of 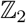 and a *ρ*-feature space *F* with *G*-group action *π*, we say that a pointwise extension 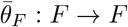 is equivariant if it commutes with *π*, i.e., 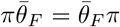. By definition of the group action (1), this is equivalent to saying that the pointwise extension 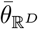 of *θ* to 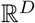 commutes with *ρ*. The following theorem gives an exhaustive characterization of a large class of equivariant pointwise extensions for any *ρ*-feature space:

#### Theorem 3.

*Let ρ be a representation of* 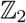 *and* 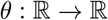 *be a continuous function with left and right derivatives at* 0. *Let F be a ρ-layer and* 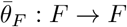 *be the point-wise extension of θ on this layer. Then* 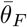 *is equivariant if and only if at least one of the following cases holds*:

1. *θ is a linear function*.
2. *θ is an affine function, and ρ*(−1)1 = 1.
3. *θ is not an affine function, and there exists a permutation matrix* Π, *integers* 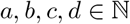, *and scalars* 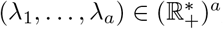, *such that ρ decomposes as*

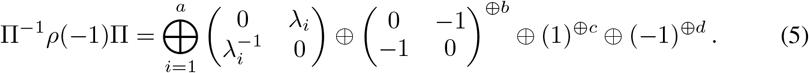 *In that case*,
  - *Either b* = *d* = 0 *and* ∀*i*, λ_*i*_ = 1 *and θ is any function*,
  - *Or b* = *d* = 0 *and* ∃*i*, λ_*i*_ ≠ 1 *and θ is a leaky ReLu function*.^2^
  - *Or b* + *d* > 0 *and* ∀*i*, λ_*i*_ = 1 *and θ is an odd function*,

The first case in Theorem 3 is of little interest, since pointwise linear functions are always equivariant to linear group actions. The second case essentially says that adding a constant to a pointwise linear function is only equivariant for representations *ρ* such that the sum of all rows of *ρ*(−1) is equal to 1. This holds for example for the regular representation and the RCPS layers, but not for an irrep feature space of type (*I, a, b*) with *b* > 0, since in that case, some rows have a single “-1” entry. The most interesting case is the third one, since it describes what pointwise nonlinearities one can apply. The condition (5) on the decomposition of *ρ* essentially excludes all representations that have more than one nonzero value in at least one row of *ρ*(−1). Among valid *ρ*’s that decompose as (5), we see that the regular representation (corresponding to the first block in (5) with λ_*i*_ = 1)), used in RCPS, stands out as the only that allows *any* nonlinearity, besides of course invariant channels of type “+1” (third block in (5)). Replacing a “1” in the regular representation by a scalar λ_*i*_ ≠ 1 (in the first block of (5), with *b* = *d* = 0) creates a valid representation *ρ*, however only leaky ReLu pointwise nonlinearities can be applied in that case. Another case of practical interest is the irrep feature space of type (*I, c, d*) for some *c* > 0 and *d* > 0. By Theorem 3, only odd nonlinearities are allowed in that case, such as the hyperbolic tangent function. Finally, one should keep in mind that other representations, which do not satisfy the conditions listed in Theorem 3, do not allow any equivariant nonlinear pointwise transform; this is for example the case of 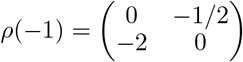, which is a valid representation of 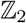 but does neither meet the condition to accept affine activations (case 2), nor to accept nonlinear activations (case 3) because *ρ*(−1) does not decompose according to (5).

#### Other activation functions

Besides pointwise transformations from a *ρ*-feature space to itself characterized in Theorem 3, the set of nonlinear equivariant layer is tremendous and the design choices are endless. A first extension is to keep pointwise activation, but to allow different nonlinearities on different channels, e.g., by using any function on the “+1” channels and an odd function on the“-1” channels of an irrep feature space. Another relaxation is to use different input and output representations. While odd functions will not affect the field type, even functions will turn a field of type “-1” into a “+1” type. It is well known that any function decomposes into a sum of an odd and even function. Therefore, given *ρ*, a representation decomposed as in (5), any point-wise non-linearity can be used in a *ρ*-feature space by first decomposing it into its odd and even components and applying each component separately for the second and fourth blocks.

Other possibilities exist and include creating new representations by tensorization, which amounts to taking pointwise products between different channels [13, 21, 37]. or using non point-wise activation layers, that act on several coupled dimensions, such as the ones used in [37]. For instance, we could apply the max function to paired channels. These possibilities are discussed in [39]

#### Batch normalization

An equivariant batch normalization was introduced by [34]. It considers a feature map and its reverse complement as two instances, which is easy to do because the reverse complement feature map is already computed when using regular representation. We propose another batch normalization for irrep feature spaces that also gives the result we would have had if the batch contained all the reverse complement of its sequences. For the “+1” dimensions, it amounts to scaling as we would have the same values twice. For the “-1” dimensions, we enforce a zero mean and compute a variance estimate based on this constraint.

#### K-mers

Instead of the standard one-hot encoding of individual nucleotides as input layer, we propose to one-hot encode *k*-mers for *k* ≥ 1, i.e., overlapping blocks of *k* consecutive nucleotides. This technique is known to improve performance in several tasks [27, 28]. In order to implement it into an equivariant network, we need to know how the group acts on the *k*-mers space, made of 4^*k*^ elements. The simplest idea is to pair the index of the channels of two RC *k*-mers. Because some *k*-mers are their own reverse complement, the canonical way to do so is to have a representation that is a blend of “+1” irrep and regular representation. An alternative is to make the regular representation act on the *k*-mers instead by redundantly encoding these *k*-mers into paired dimensions. This is the strategy we follow in our implementation, to be more coherent with the usual input group action.

## 3 Experiments

We assess the performance of various equivariant architectures on a set of three binary prediction and four sequence prediction problems used by Zhou et al. [45] to assess the performance of RCPS networks. The binary classification problems aim to predict if a DNA sequence binds to three transcription factors (TFs), based on genome-wide binarized TF-ChIP-seq data for Max, Ctcf and Spi1 in the GM12878 lymphoblastoid cell-line [34]. The sequence prediction problems aim to predict TF binding at the base-pair resolution, using genome-wide ChIP-nexus profiles of four TFs-Oct4, Sox2, Nanog and Klf4 in mouse embryonic stem cells. For a more detailed explanation of the experimental setup, please refer to Zhou et al. [45]. We report “significant” differences in performance below when the P-value of a Wilcoxon signed rank test is smaller than 0.05.

### Models

We build over the work of Zhou et al. [45] for both the binary and the sequence prediction problems. They benchmarked an equivariant RCPS architecture and a corresponding non-equivariant model, with the same number of filters and trained with data augmentation, which we respectively refer to as “RCPS” and “Standard” models below. The data augmentation scheme for the “Standard” model consists in adding to the training set the reverse complement sequences of all training sequences, which is a natural procedure to let the model “learn” the equivariance without encoding it explicitly in the architecture of the network. We checked empirically that data augmentation significantly improves the performance of non-equivariant models (Appendix A.6.1). In addition, we extend the RCPS architecture with one-hot encoding of *k*-mers as input layers, which we refer to as “Regular” below. Finally, we add to the comparison a new equivariant network where each RCPS layer is replaced by an (*I, a, b*) layer with the same number of filters, which we call “Irrep” below. We also use *k*-mers and vary the ratio *a*/(*a* + *b*) in this model. We combine the regular and “+1” dimensions with *ReLu* activations and the “-1” dimensions with a *tanh* activation.

### Influence of hyperparameters in equivariant models

To assess the impact of different hyperparameters in the family of equivariant models we propose (*k*-mer length for Irrep and Regular, *a*/(*a* + *b*) ratio for Irrep), we train equivariant models with different combinations of hyperparameters on the training set and assess their performance on the validation set, repeating the process ten times with different random seeds. We assess the performance of each run in terms of Area under the Receiver Operator Characteristic (AuROC), and show in Figure 2 the average performance reached by all runs with a given ratio *a*/(*a* + *b*) ∈ {0, 1/4, 1/2, 3/4, 1} (left) and with a given *k* ∈ {1, 2, 3, 4} (right). We see a clear asymmetry in the performance as a function of *a*/(*a* + *b*), with poor performance when *a* = 0 and optimal performance for *a* = 0.75, significantly better than all other ratios tested. This confirms that exploring different irreps may be valuable. As for the *k*-mer length, setting *k* = 3 gives the best performance and significantly outperforms all other values of *k* tested. This confirms that going beyond one-hot encoding of nucleotides in equivariant architectures can be beneficial.

**Figure 2:**
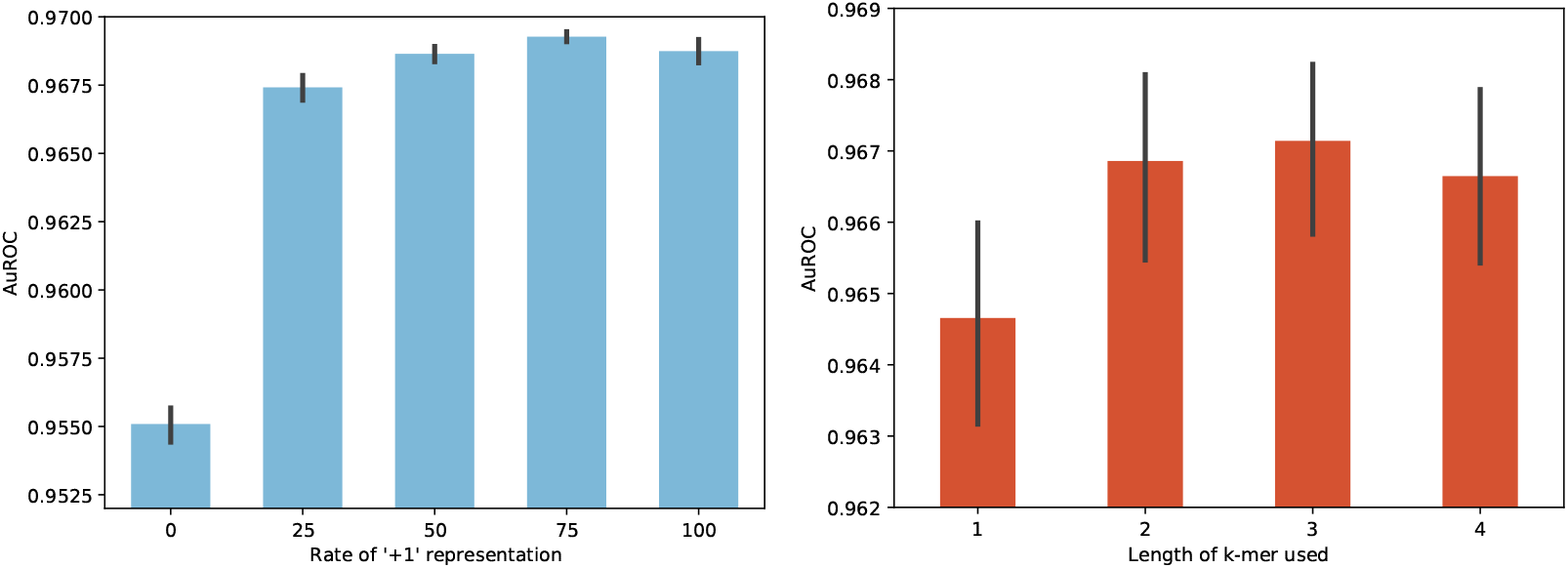
Average AuROC performance across four TFs and 10 random seeds for the Irrep model as a function of *a*/(*a*, + *b*) (*left*, also averaged over *k* values) and for the Irrep and Regular models as a function of *k* (*right*, also averaged over *a*/(*a* + *b*) values for Irrep).

### Binary task

We then compare the test set performance of three different models for the binary classification task: 1) Standard, 2) RCPS, and 3) the best Irrep or Regular equivariant model, where hyperparameters are selected based on the AuROC on the validation set, which we denote as “Best Equivariant”. Figure 3 (left) shows the performance of each model on each TF task and overall. As already observed by [34], the equivariant RCPS architecture has a strong lead over the Standard, non-equivariant model in spite of data augmentation. Interestingly, we see that Best Equivariant is significantly better than RCPS on all tasks, and that the performance gain from RCPS to Best equivariant is of the same order as the performance gain from Standard to RCPS. This demonstrates that the family of equivariant architectures we introduce in this paper can lead to significant improvement over existing architectures.

**Figure 3:**
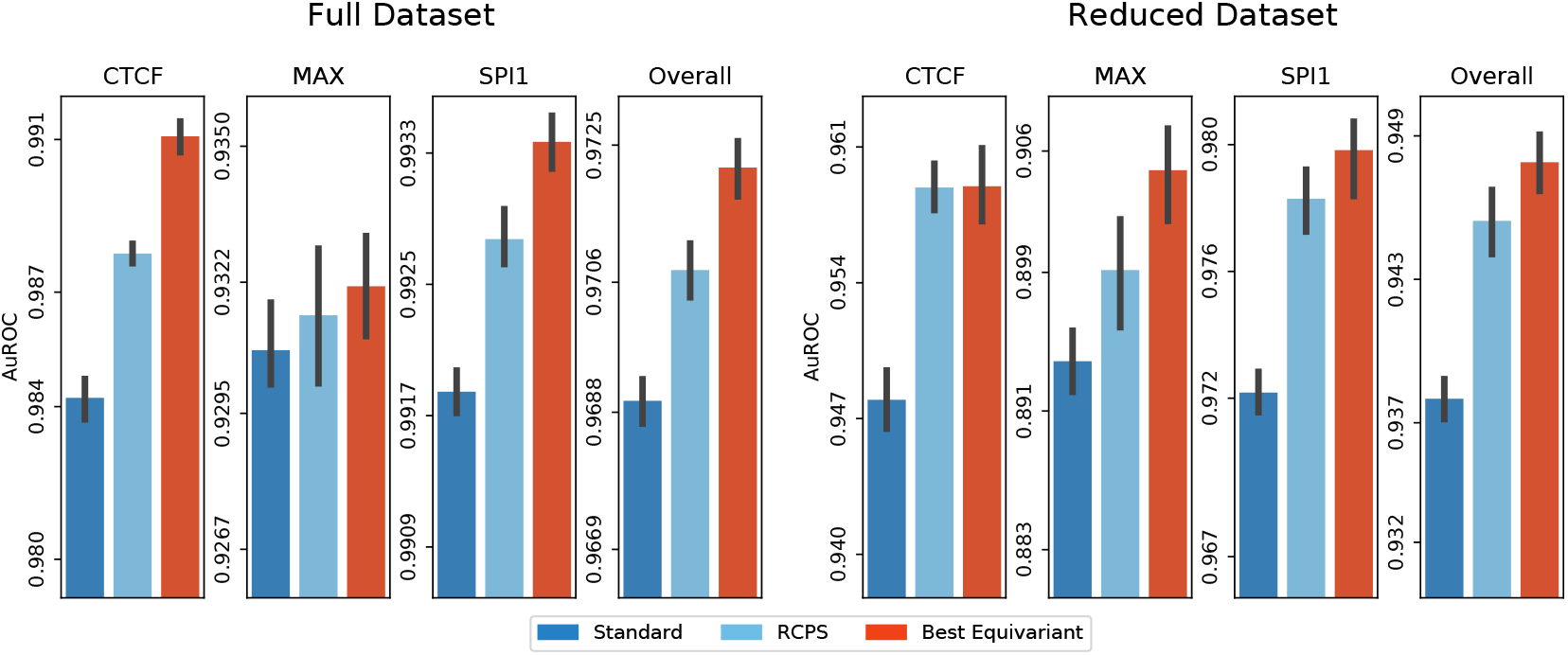
AuROC performance of the three different models (Standard, RCPS and Best equivariant after hyperparameter selection on the validation set) on the three binary classification problems CTCF, MAX and SPI1, as well as their average. Error bars correspond to an estimate of the standard error on 10 repeats with different random seeds. The left plot is the performance on the full datasets, while the right plot shows the performance where models are trained on a subset of 1,000 sequences only (notice the differences of AuROC values on the vertical axis in both plots).

### Reduced models

Since equivariant architectures are meant to be particularly beneficial in the low-data regime [15], we further assess the performance of the three models on the same binary classification problems but with only 1,000 sequences used to train the models, and show the results on Figure 3 (right). Overall, the performances are worse than in the full-data regime (Figure 3, left), which confirms that this is a regime where more data helps. We also see that the relative order of the three different methods remains overall the same, with Best Equivariant outperforming RCPS, which itself outperforms Standard. Interestingly, the gaps between the best and worse models widens in the low-data regime, showing that the prior is more useful in this setting. More precisely, there is a large gap of about 1% between Best Equivariant and Standard in the low data regime, compared to a gap of about 0.3% on the full dataset. We also investigated whether equivariant models converge faster to their solutions, but found not noticeable difference (Appendix A.6.2).

### On post-hoc models

Zhou et al. [45] introduced the so-called *post-hoc* model, another equivariant method obtained by averaging the predictions of a Standard model over a sequence and its reversecomplement, and showed that it is competitive with and often outperforms RCPS. The post-hoc model only requires training and storing one network, but aggregates two predictions for each sequence at inference time. Because of that, the good performance of post-hoc may be due in part to the aggregation step common to all ensemble models [12]. To decipher the respective contributions of the network architecture, on the one hand, and of the aggregation of predictions, on the other hand, we add to the comparison an ensemble of two Standard models trained with different random seeds (*Ensemble Standard*) and an ensemble of two equivariant Irrep models (*Ensemble Irrep*) and present the results in Figure 4. We see that Ensemble Irrep strongly outperforms Best Equivariant, and both post-hoc and Ensemble Standard widely outperform the Standard architecture. This confirms that ensembling equivariant or non-equivariant models through post-hoc of ensemble aggregation is always useful (at the cost of increased computational time). We see that Ensemble Standard is not significantly different from post-hoc Standard on CTCF and SPI1, but that post-hoc Standard is better on MAX, suggesting that most of the benefits of post-hoc Standard indeed comes from the ensembling effect. Regarding the impact of the architecture for a given budget of predictions, we saw earlier than Best equivariant significantly outperforms Standard when a single prediction per test sequence is allowed, and see now that Ensemble Irrep strongly outperforms both post-hoc and Ensemble Standard when two predictions are allowed, thus confirming the benefit of equivariant architectures in all settings. We also see that a single Best equivariant models outperforms post-hoc and Ensemble Standard, indicating that enforcing equivariance throughout the network is not only faster but also more more accurate than averaging a non-equivariant model over group transformed inputs.

**Figure 4:**
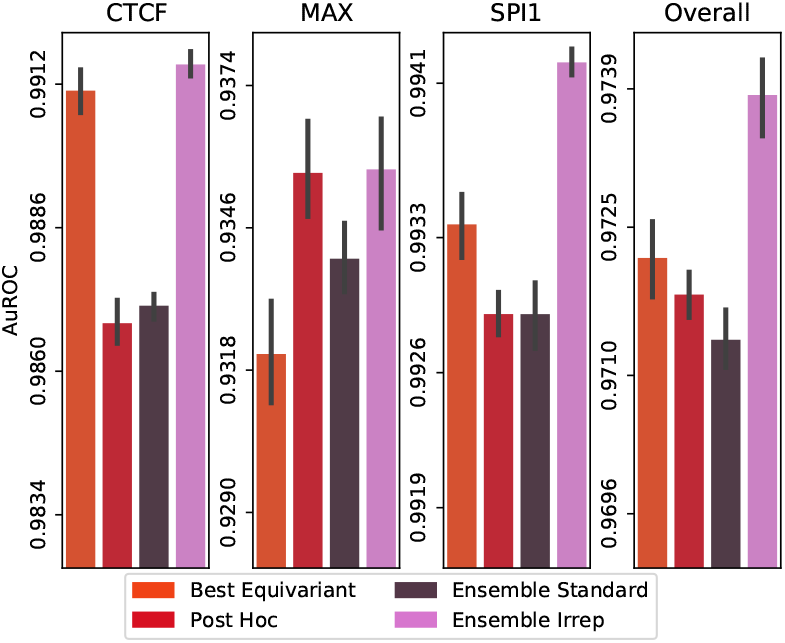
AuROC performance on the three binary classification problems, for the Best Equivariant model, the post-hoc Standard model, and an ensemble of two Standard or Irrep models. Error bars correspond to an estimate of the standard error on 10 repeats with different random seeds.

### Profile task

We now compare the performance of different models on the profile prediction tasks. To limit the carbon footprint of this study, and based on the influence of hyperparameters on the binary task (Figure 2), we only test two equivariant models in addition to Standard and RCPS: a Regular model with *k* = 3, and an Irrep model with *k* = 3 and *a*/(*a* + *b*) = 75%. We also assess the performance of post-hoc Standard (the best model in [45]), and an ensemble of two models of the best performing equivariant model. Figure 5 shows the performance of all models in terms of Spearman correlation between the target profile and the predicted ones, on the full dataset (left) or a reduced experiment with only 1,000 training sequences (right).

**Figure 5:**
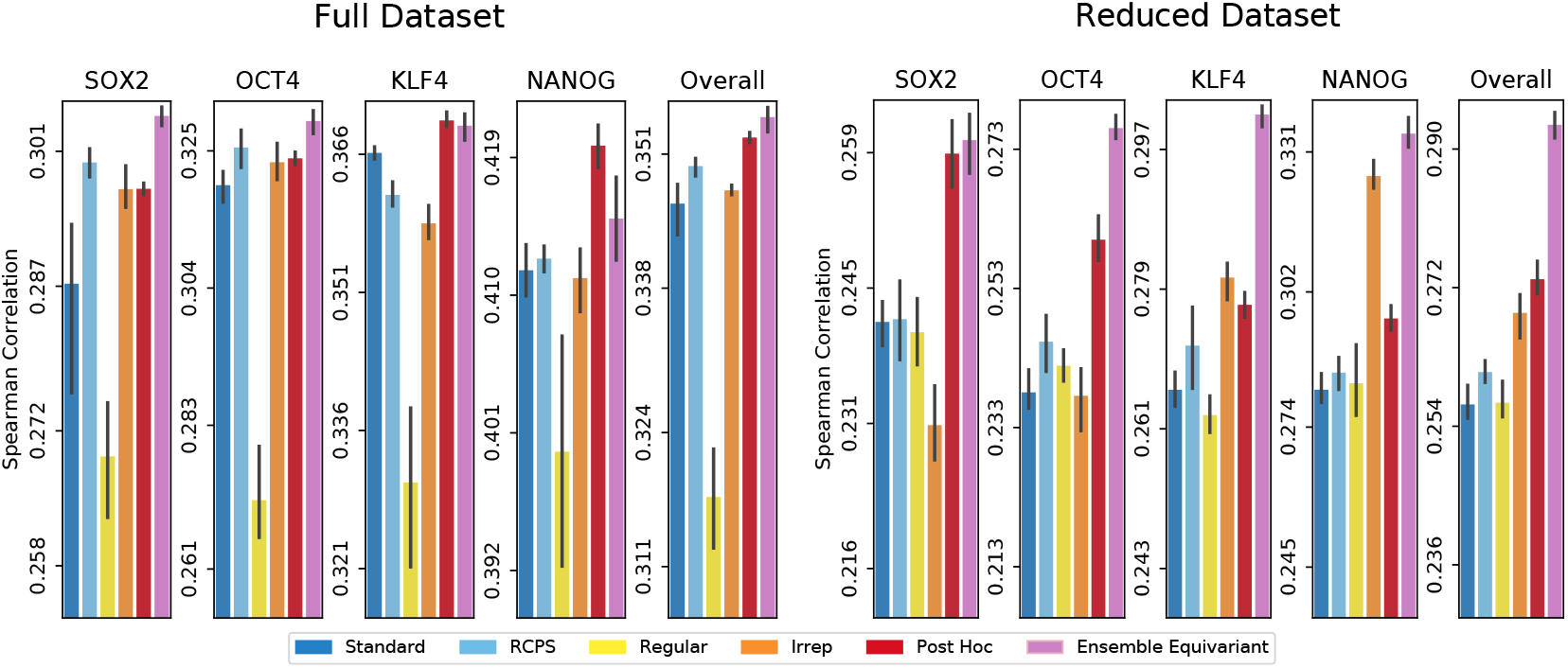
Spearman Correlation between true and predicted profiles by different methods for four data sets.

First of all, we observe as before that in the low-data regime, the gap between standard and equivariant networks grows in favor of equivariant ones. We also observe, surprisingly, that Irrep, which outperformed RCPS on the binary task, now underperforms it. A possible explanation could be that since this task aims to annotate an individual nucleotide, encoding the nucleotide level information using *k*-mers makes the signal blurry and decreases performance. However, in the reduced setting, Irrep performs better again. These results indicate that for now the best model should be chosen empirically on a validation set. Finally, despite good performance of post-hoc Standard, the ensemble equivariant model once again performs better for the same computational cost at inference.

### Experiment settings and computational cost

All experiments were run on a single GPU (either a GTX1080 or a RTX6000), with 20 CPU cores. The binary classification experiments were shorter to train. To limit our carbon footprint, we chose to run more experiments on this task, e.g., for hyperparameter tuning and to reduce the number of replicates for the profile task. The total runtimes of each of those tasks were approximately of a week.

## 4 Conclusion

In this paper, we addressed the problem of including the RC symmetry prior in neural networks. Leveraging the framework of equivariant networks, in particular steerable CNNs, we deepened existing methods by unraveling the whole space of linear layers and pointwise nonlinearities that are translation and RC-equivariant. We also investigated the links between the linear representations and the non-linear layers of neural networks, exposing the special role of the regular representation in equivariant networks. Finally, we implemented new linear and nonlinear equivariant layers and make all these equivariant layers available in Keras [5] and Pytorch [31].^3^ We then explored empirically how this larger equivariant functional space behaves in terms of learning. Our best results improve the state of the art performance of equivariant networks, showing that new equivariant architectures can have practical benefits. In the future we plan to test more deeply the newly proposed architectures on prediction tasks involving double-stranded DNA, such as DNA-protein binding prediction, epigenetics or metagenomics. On the theoretical side, we characterized equivariant pointwise nonlinearities that preserve the layer type, but more general nonlinear transforms (e.g., not pointwise, or changing the layer type) remain to be fully characterized.

## Acknowledgments and Disclosure of Funding

V.M. is recipient of a doctoral fellowship from the INCEPTION project [PIA/ANR-16-CONV-0005] and benefits from support from the CRI through Ecole Doctorale FIRE - Programme Bettencourt. We thank Marie Dechelle, Jacques Boitreaud, Carlos G. Oliver and Guillaume Bouvier for reviewing the manuscript. We thank Avanti Shrikumar, Hannah Zhou and Anshul Kundaje for helpful discussions and sharing their code.

## A Appendix

### A.1 Illustration of group actions

This section is intended to provide a visual, more intuitive understanding of the different group actions on the tensors of our network. We begin with a visualization of the group action for the input space. We exemplify it over the sequence `GGACT`, whose reverse complement is `AGTCC`. The sequence is one hot encoded as explained in the main text and the group action over 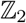 consist in flipping the tensor along the spatial axis and swapping the channels pairwise.

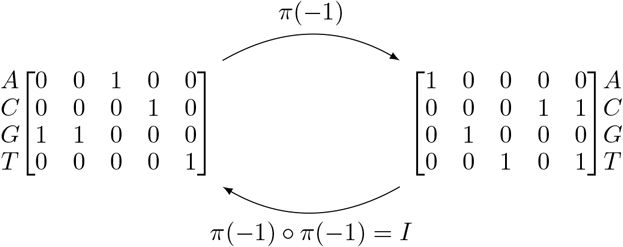

Now we illustrate the actions of other representations, on an example tensor 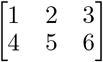 with two channels (of type *a* or *b*) and three positions; this could typically be the representation of an input sequence of length 3 in an intermediate layer of dimention 2. Choosing the canonical representations of type (*I*, 2, 0), (*I*, 0, 2) and (*I*, 1, 1) respectively, we get the following group actions (for clarity we add the channel type, *a* or *b*, near each matrix row):

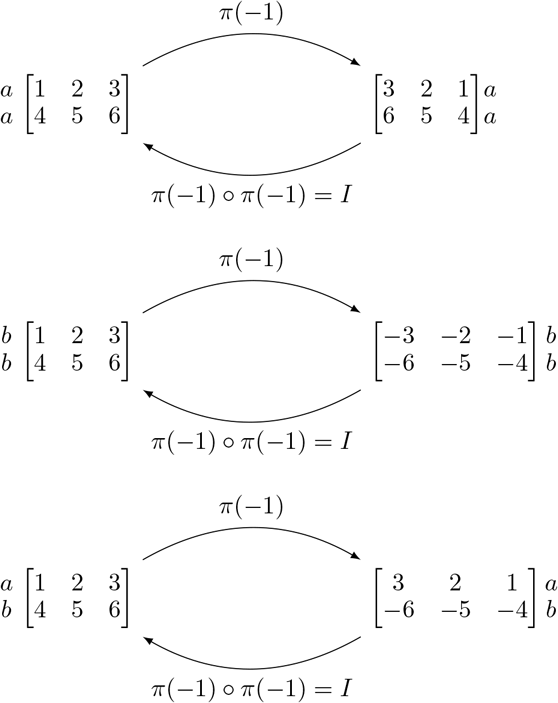

Finally, when using different values for P, we can get other group actions. As mentioned in the main text, by choosing (*P_reg_*, 1, 1), where 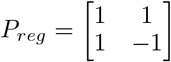, we get the regular representation that flips the input channel. We also provide an example of the group action for a general P matrix, by choosing (*P_general_*, 1, 1), where 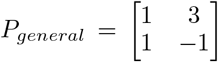, we get a representation on the fibers 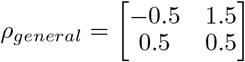

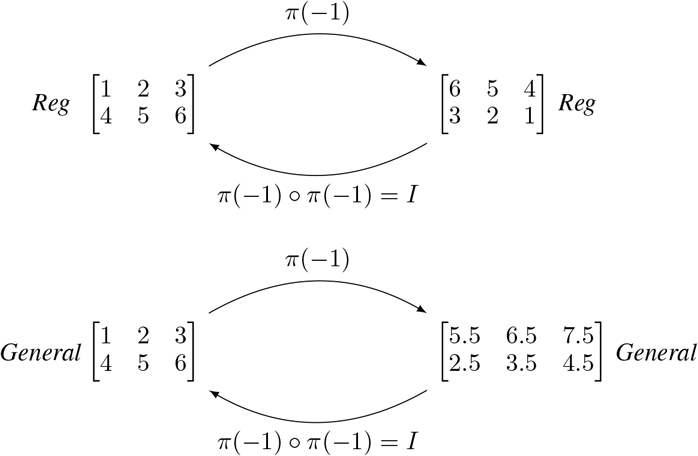

Over the course of these examples, we have limited ourselves to the case where the input tensor had only three nucleotides and two channels, but this is coincidental. The representation with arbitrary P can mix an arbitrary number of channels together with the group action.

### A.2 Proof of Theorem 1

#### Proof.

The irreducible representations (irreps) of the 2-elements group 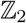 are the 1-dimensional trivial and sign representations, given respectively by *ρ*_1_(*s*) = 1 and *ρ*_1_(*s*) = *s*. Any representation *ρ_n_* can be decomposed as a direct sum of irreps, and since each irrep is 1-dimensional this means that there exists an invertible matrix *P* such that *P_ρ_n__*(*s*)*P*^−1^ is diagonal, with diagonal terms either equal to 1 or equal to s. If we denote by *a_n_* (resp. *b_n_*) the number of diagonal terms equal to 1 (resp. s), then Theorem 1 follows.

### A.3 Proof of Theorem 2

#### Proof.

Cohen et al. [11, Theorem 3.3] gives a general result about linear equivariant mapping. We first show that this result can be applied here, to show that these linear mappings are exactly the ones written as (2) and (3). For sake of clarity, we then provide a fully self-contained proof of the same result.

Let us first show that (2) and (3) correspond to a particular case of Cohen et al. [11, Theorem 3.3]. Under the notations of [11], our group is 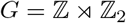, a locally compact, semi-direct product group. We choose 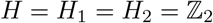, making the coset space 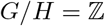. Since our group is a semi direct product group, we have *h*_1_(*x, s*) = *s*. The spaces *F_n_* that we have considered are signals in 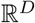 over the coset space, acted upon by the representation induced by *ρ*. Equivalently, they are sections of the associated vector bundle for the trivial case of a product group. Therefore, these *F_*n*_* exactly coincide with the setting of Cohen et al. [11, Theorem 3.3] and {*ϕ* : *F_n_* → *F*_*n*+1_|*π*_*n*+1_*ϕ* = *ϕπ_n_*} is exactly 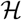. Then, by [11, Theorem 3.3], *ϕ*: *F_n_* → *F*_*n*+1_ is equivariant if and only if it can be written as a convolution:

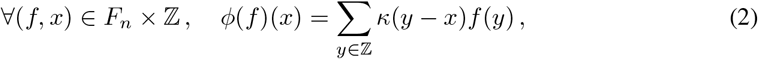

where the kernel 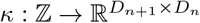 satisfies:

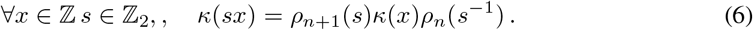

Using that for 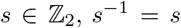, *s*^−1^ = *s*, and the triviality of this equation for *s* = 1, we get that (6) is equivalent to (3)

For sake of clarity and completeness, we now provide a more explicit and self-contained proof for (2) and (3), that follows the one of [40, Theorem 2] in our specific setting. We first notice that any linear mapping *ϕ*; *F_n_* → *F*_*n*+1_ can be written as

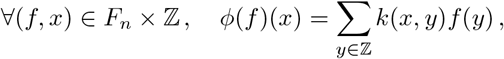

for some function 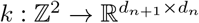. For any *g* = *ts* ∈ *G*, the action of *G* on *F*_*n*+1_ gives:

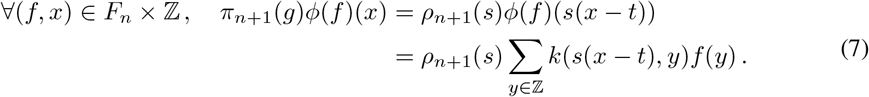

Similarly, the action of *G* on *F_n_* followed by *ϕ* gives:

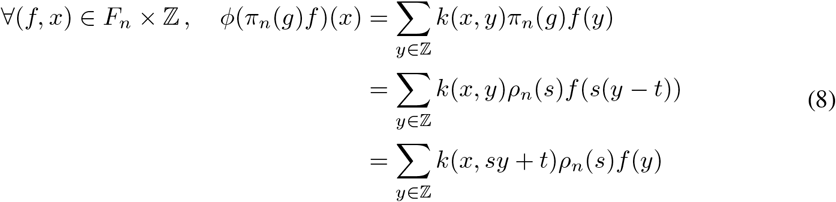

where we made the change of variable *y* ↦ *sy* + *t* to get the last equality. *ϕ* is equivariant if and only if, for any *g* ∈ *G*, *ϕ* ○ *π_n_*(*g*) = *π*_*n*+1_(*g*) ○ *ϕ*, which from (7) and (8) is equivalent to:

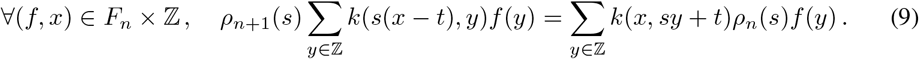

For any 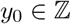 and 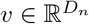, let us apply this equality to the function *f* ∈ *F_n_* given by *f*(*y*_0_) = *v* and *f*(*y*) = 0 for *y* ≠ *y*_0_:

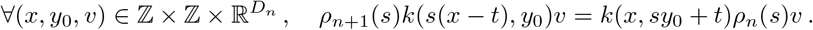

Since this must hold for any 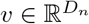 this necessarily implies:

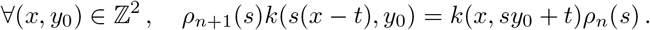

With the change of variable *y* = *s*(*y*_0_ − *t*), this is equivalent to:

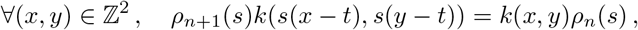

which itself is equivalent to

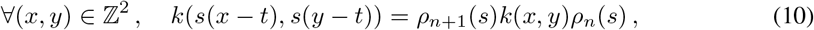

where we used the fact that *ρ*_*n*+1_(*s*)^2^ = *ρ*_*n*+1_(*s*^2^) = *I* for any 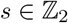. This must hold in particular for *s* = 1 and *t* = *x*, which gives:

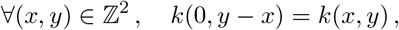

i.e., *k* is necessarily translation invariant in the sense that there must exist a function 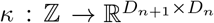 such that

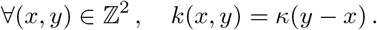

From (10) we see that *κ* must satisfy

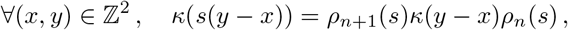

which boils down to the following constraint, after observing that the constraint is always true for *s* = 1 and is therefore only nontrivial for *s* = −1:

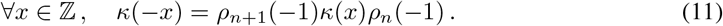

At this point, we have therefore shown that an equivariant linear function must have an expansion of the form

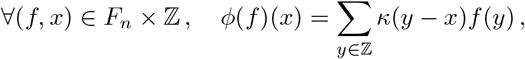

where *κ* must satisfy (11). Conversely, such a linear layer trivially satisfies (9), and is therefore equivariant. This proves (2) and (3).

To prove (4), we simply rewrite (3) using Theorem 1:

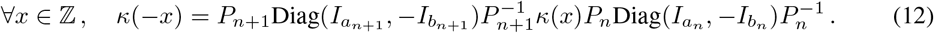

Thus writing the matrix 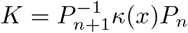 by blocs of sizes *a*_*n*+1_ × *a_n_*, *a*_*n*+1_ × *b_n_*, *b*_*n*+1_ × *a_n_* and *b*_*n*+1_ × *b_n_*, we have :

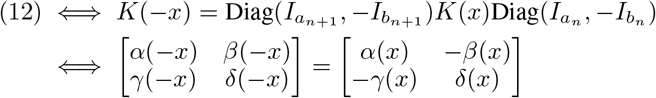

This gives us the equivalence (3) ⇔ (12) ⇔ (4).

### A.4 Resolution of the constraint for other basis

To go from an arbitrary representation (*P, a, b*) to another, we can write an odd/even kernel and change of basis. One may also solve the constraints (3) for specific representations, and save the need of multiplication by *P*_*n*+1_ and 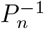 in (4). In this section, we solve the constraint in other basis, to go from one kind of representation (irrep or regular) to another. We just substitute the correct representation and see what constrained kernel it gives. The irrep and regular representations are in a basis such that they write as :

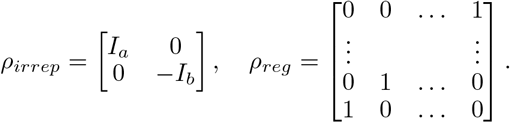

We get the following table of constraints :

| $F_n \backslash F_{n+1}$ | 'irrep' | 'regular' |
| --- | --- | --- |
| 'irrep' | $\begin{bmatrix} \alpha(-x) & \beta(-x) \\ \gamma(-x) & \delta(-x) \end{bmatrix} = \begin{bmatrix} \alpha(x) & -\beta(x) \\ -\gamma(x) & \delta(x) \end{bmatrix}$ | $[\kappa_{j,a}(-x), \kappa_{j,b}(-x)] = [\kappa_{n-j,a}(x), -\kappa_{n-j,b}(x)]$ |
| 'regular' | $\begin{bmatrix} \kappa_{a,j}(-x) \\ \kappa_{b,j}(-x) \end{bmatrix} = \begin{bmatrix} \kappa_{a,n-j}(x) \\ -\kappa_{b,n-j}(x) \end{bmatrix}$ | $\kappa_{i,j}(-x) = -\kappa_{n-i,n-j}(x)$ [34] |

### A.5 Proof of Theorem 3

With a slight abuse of notations, in this section we denote the matrix *ρ*(−1) simply by 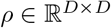, and for any 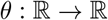 we define 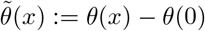. We start with three technical lemmas, before proving Theorem 3.

#### Lemma 4.

*Let h* : 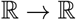 *be a continuous function with left and right derivatives at 0. If there exists* 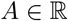 *with* |*A*| > 1 *such that*

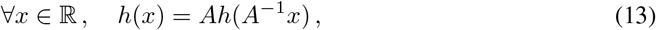

*then h is a leaky ReLu function, i.e., there exists* 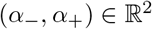 *such that*

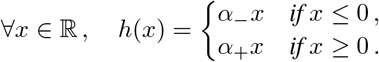

*In addition, if A* < −1, *then α*_−_ = *α*_+_, *i.e., h is linear*.

*Proof*. Equation (13) implies *h*(0) = 0 and

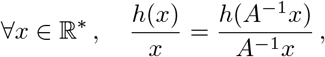

which by simple induction gives more generally:

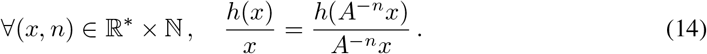

The right-hand side of (14) for *n* = 2*k* converges to 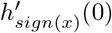 when *k* → +∞, which by unicity of the limit must be equal to the left-hand side. As a result, for any 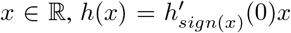, i.e., *h* is a leaky ReLu function with 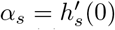 for *s* ∈ {−, +}. If in addition *A* < −1, then (14) for *n* = 2*k* + 1 converges to 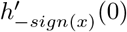 when *k* → +∞. By unicity of the limit, this implies 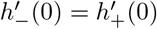, i.e., *α*_−_ = *α*_+_.

#### Lemma 5.

*Under the assumptions of Theorem 3, if* 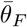 *is equivariant and if there exists* (*i, j*) ∈ [1, *D*]^2^ *such that ρ_ij_* ∉ {−1, 0, 1}, *then necessarily* 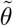 *is a leaky ReLu function*.

*Proof*. For any (*i, j*), applying the equivariance constraint *θ*(*ρx*)_*i*_ = *ρθ*(*x*)_*i*_ to the vector *x* = *ae_j_*, for any 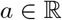, gives the equation:

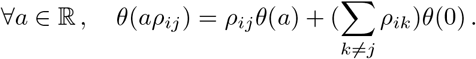

If |*ρ_ij_*| > 1, we can rewrite it as

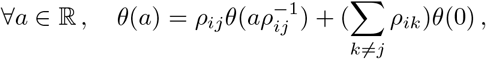

and if 0 < |*ρ_ij_*| < 1 we can rewrite it as

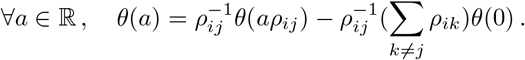

In both cases, this is an equation of the form

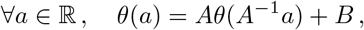

where |*A*| > 1. Subtracting to this equation the same equation written for *a* = 0 gives

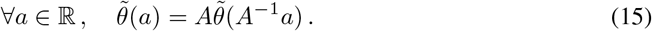

By Lemma 4, 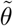 is a leaky ReLu function.

#### Lemma 6.

*Under the assumptions of Theorem 3, if* 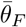 *is equivariant and if there exists at least one row in ρ with at least two nonzero entry, then necessarily θ is an affine function*.

*Proof*. Let us suppose that *ρ* contains at least a row *i* with two nonzero entries, say *ρ_ij_* ≠ 0 and *ρ_ik_* ≠ 0. Then taking *x* = *x_j_e_j_* + *x_k_e_k_* with *x_j_*, 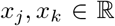, the equivariance constraint for the *i*-th dimension gives

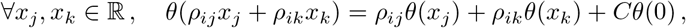

with *C* = Σ_*p*∉{*j,k*}_ *ρ_ip_*. Subtracting to this equation the same equation written for *x_j_* = *x_k_* = 0 allows us to remove the constant term and get

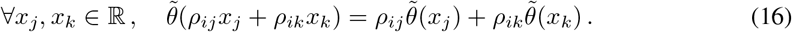

We now prove that 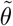 is necessarily a leaky ReLu function, i.e., that there exist 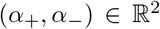 such that 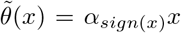, with potentially *α*_+_ ≠ *α*_−_. By Lemma 5 this is true if |*ρ_ij_*| ≠ 1 or |*ρ_ik_*| ≠ 1, so we focus on the case |*ρ_ij_*| = |*ρ_ik_*| = 1, which we decompose in two subcases. First, if *ρ_ij_* = *ρ_ik_* = *s* with *s* ∈ {−1, 1}, then taking *x_j_* = *x_k_* = *a* in (16) gives 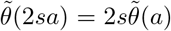, for any 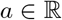. Second, if *ρ_ij_* = −*ρ_ik_* = 1 (resp. *ρ_ij_* = −*ρ_ik_* = 1), then taking *x_j_* = 2*a* and *x_k_* = *a* (resp. *x_j_* = *a* and *x_k_* = 2*a*) gives 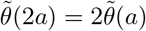. In both subcases, by Lemma 4, 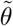 must be a leaky ReLu function.

Knowing that 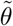 is a leaky ReLu function with coefficients *α*_+_ and *α*_−_, in order to prove that *θ* is necessarily an affine function (i.e., that 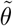 is linear), we need to show that *α*_+_ = *α*_−_. For that purpose, let us first suppose that *ρ_ij_* and *ρ_ik_* are both positive or both negative. Then there exists a pair 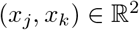 such that *x_j_* > 0, *x_k_* < 0 and *ρ_ij_x_j_* + *ρ_ik_x_k_* < 0. Similarly, if *ρ_ij_* and *ρ_ik_* are of different signs, say without loss of generality *ρ_ij_* < 0 and *ρ_ik_* > 0, then any pair 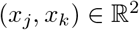 such that *x_j_* > 0, *x_k_* < 0 satisfies *ρ_ij_x_j_* + *ρ_ik_x_k_* < 0. In both cases, using the fact that 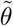 is linear on 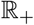 and on 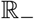, (16) gives

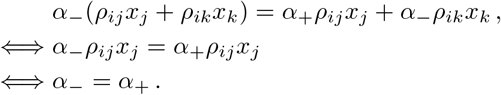

We are now ready to prove Theorem 3.

#### Proof of Theorem 3.

To characterize the functions *θ* and representations *ρ* such that 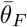 is equivariant, we proceed by a disjunction of cases on *θ*, depending on whether it is affine.

If *θ* is affine, say *θ*(*x*) = *αx* + *β*, then 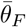 is equivariant if and only if, for any 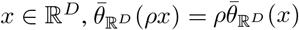. This is equivalent to

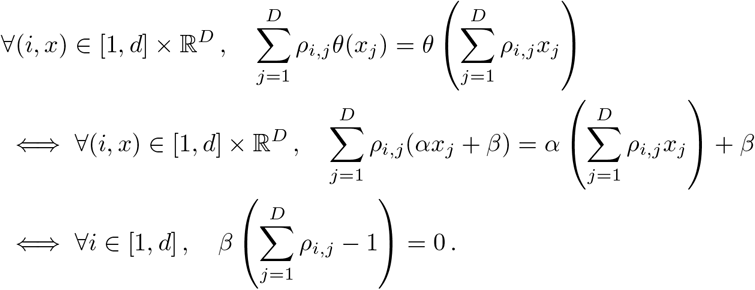

This shows that if *θ* is affine, then 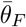 is equivariant if and only *β* = 0, i.e., *θ* is linear (case 1 of Theorem 3), or *ρ***1** = **1** (case 2 of Theorem 3).

If *θ* is not affine and 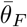 is equivariant, then by Lemma 6 we know that *ρ* can have at most one nonzero entry per row. Since *ρ* is invertible, it must have at least one nonzero entry per row, so we conclude that if contains exactly one nonzero entry per row, hence a total of *D* nonzero entries. Being invertible, it must also contain at least one nonzero entry per column, so we conclude that it contains also exactly one nonzero entry per column. Using the fact that *ρ*^2^ = *I*, we can further clarify how nonzero entries must be organized:

- For a nonzero entry *ρ_ii_* ≠ 0 on the diagonal, we must have 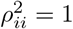, i.e., *ρ_ii_* ∈ {−1, +1}.
- For an off-diagonal nonzero entry *ρ_ij_* ≠ 0 with *i* ≠ *j*, we must have *ρ_ij_ρ_ji_* = 1, i.e., 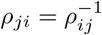.

Splitting the nonzero entries by sign, this implies that there exists a permutation matrix Π such that

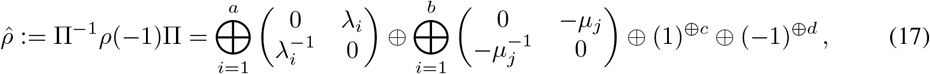

for some 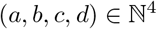 such that *a* + *b* + *c* + *d* = *D* and 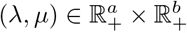. For any *i* ∈ [1, *D*], let us now denote by *τ*(*i*) the column corresponding to the nonzero entry of the *i*-th row of 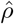, i.e., the only index such that 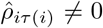. Then the action of 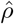 on a vector 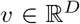 has the simple form 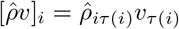. By writing the equivariance property 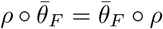 coordinate by coordinate, we can therefore say that 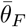 is equivariant if and only if:

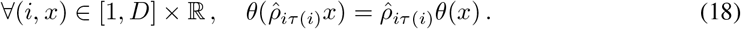

Let us now consider two possible cases:

- If there exists *i* ∈ [1, *D*] such that 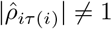, then by Lemma 5 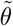 is a leaky ReLu function, i.e., there exist 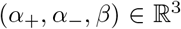 such that 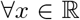, *θ*(*x*) = *α*_*sign*(*x*)_*x* + *β*. In that case, by (18), 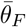 is equivariant if and only if:

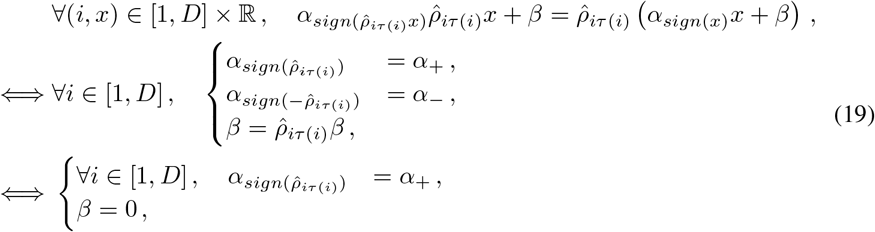

where the first equivalence comes from identifying the coefficients of the linear equation in *x* on 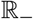 and 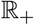, and the second equivalence comes from the observation that the two conditions in *α* in the first equivalence are themselves equivalent to each other, so we can keep only one of them, and that the condition on *β* is equivalent to *β* = 0 since we assume the existence of an *i* ∈ [1, *D*] such that 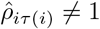. Since we assume that *θ* is not affine, we can not have *α*_−_ = *α*_+_, which by (19) rules out the possibility of having negative entries in 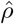, i.e., necessarily *b* = *d* = 0 in (17). If that is not the case, then the condition on *α* in (19) is automatically met for all *i* ∈ [1, *D*], so we have that 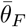 is equivariant if and only if *β* = 0, i.e., if and only if *θ* is a leaky ReLu function. This is the second statement in Case 3 of Theorem 3, when we further notice that when *b* = 0 the only entry in 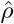 that can have been different from −1 and 1 is a λ_*i*_ in (17).
- If for all *i* ∈ [1, *D*], 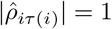, then (17) simplifies as

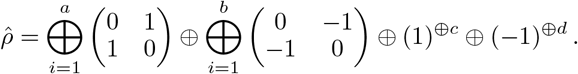 In that case, the equivariance condition (18) is particularly simple, and true for any *θ* for positive values. For each *i* such that 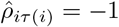 it reads 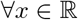, −*θ*(*x*) = *θ*(−*x*), and is therefore true if and only if *θ* is odd. Noticing that the latter constraint occurs if and only if *b* + *d* > 0 finally leads to the first and third statements in Case 3 of Theorem 3.

### A.6 Additional result

#### A.6.1 Effect of data augmentation and size for non-equivariant models

Given a non-equivariant model, a simple way to let it “learn” to be equivariant is to train it with data augmentation, where for each sequence in the training set we add its reverse complement to the training set. This doubles the size of the training set, which increases the training time. If we compare such a non-equivariant model with an equivariant model with the same number of channels in each layers, then it has about twice the same number of free parameters to train, and we therefore call it “big”; as an alternative, one may want to restrict the number of channels in each layer to enforce the same number of parameters as the equivariant model. To assess the benefits of data augmentation and number of channels, we plot in Figure 6 the performance of a standard, non-equivariant model with or without data augmentation, and with the same number of channels or half of it, on the binary classification tasks. We see that the number of channels has no significant impact on the performance, but that data augmentation has a significant positive impact. In the main text, we therefore restrict ourselves to the standard model with data augmentation as non-equivariant baseline model.

**Figure 6:**
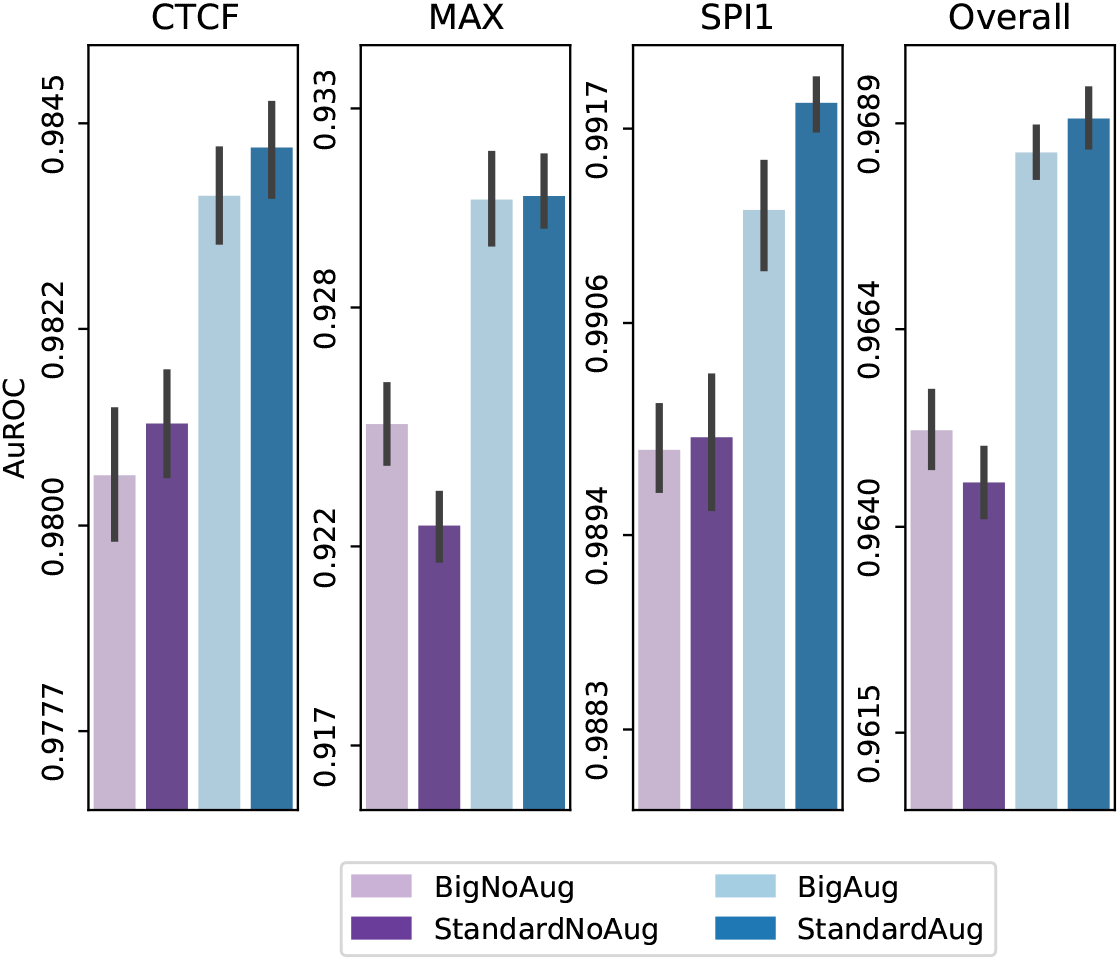
Binary task performance of a standard, non-equivariant model trained with (“Aug”) or without (“NoAug”) data augmentation, and with more (“Big”) or less (“Standard”) channels.

#### A.6.2 Comparison of learning curves

Because equivariant model are supposed to converge faster, we looked into the learning curves of our models, i.e., how the test performance increases as a function of the number of epochs during training. However, we do not see a major difference in the learning dynamics between the equivariant and non equivariant models.

**Figure 7:**
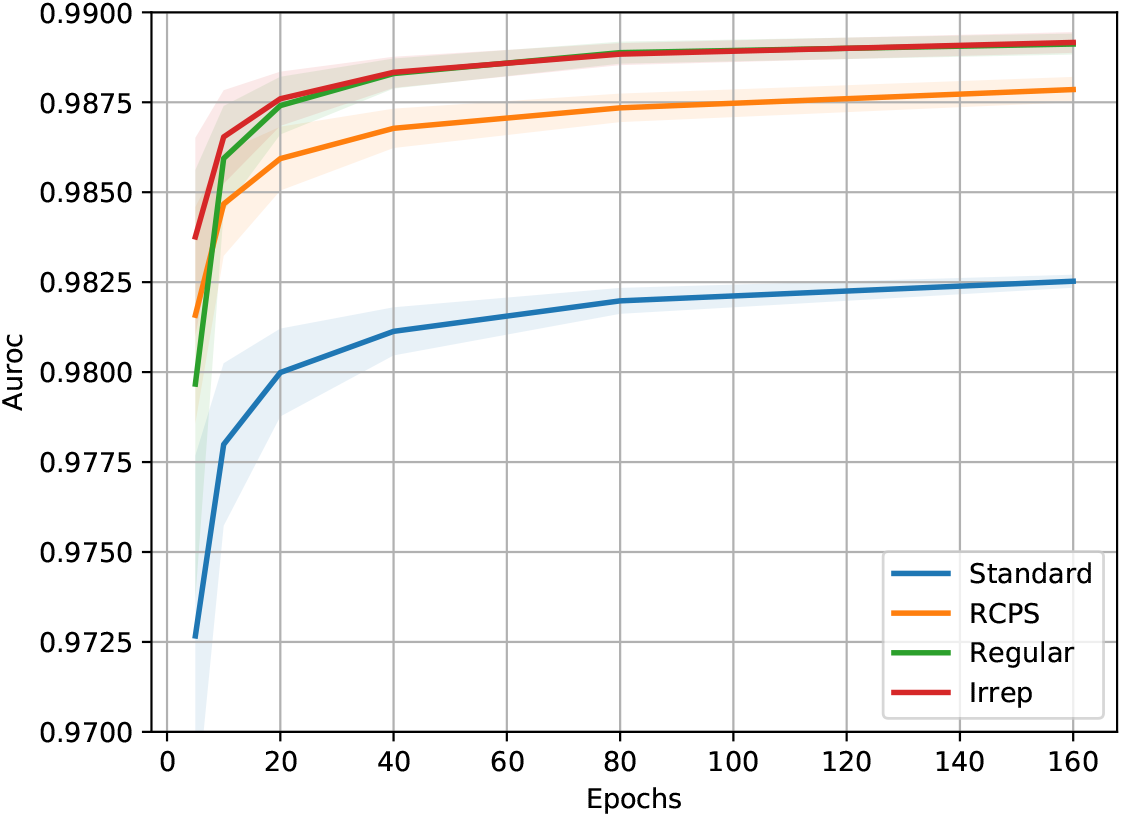
AuROC performance of the four different models on the three binary classification problems CTCF, MAX and SPI1, as well as their average over the course of learning.

## Footnotes

1 As of May, 2021: https://www.ebi.ac.uk/ena

2 A leaky ReLu function is *θ*(*x*) =*α*_*sign*(*x*)_*x* for some 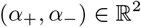.

3 code available at https://github.com/Vincentx15/Equi-RC

